# Beware the Jaccard: the choice of metric is important and non-trivial in genomic colocalisation analysis

**DOI:** 10.1101/479253

**Authors:** Stefania Salvatore, Knut Dagestad Rand, Ivar Grytten, Egil Ferkingstad, Diana Domanska, Lars Holden, Marius Gheorghe, Anthony Mathelier, Ingrid Glad, Geir Kjetil Sandve

## Abstract

**Background:** The generation and systematic collection of genome-wide data is ever-increasing. This vast amount of data has enabled researchers to study relations between a variety of genomic and epigenomic features, including genetic variation, gene regulation, and phenotypic traits. Such relations are typically investigated by comparatively assessing genomic co-occurrence. Technically, this corresponds to assessing the similarity of pairs of genome-wide binary vectors. A variety of metrics have been proposed for this problem in other fields like ecology. However, while several of these metrics have been employed for assessing genomic co-occurrence, their appropriateness for the genomic setting has never been investigated.

**Results:** We show that the choice of metric may strongly influence results and propose two alternative modelling assumptions that can be used to guide this choice. On both simulated and real genomic data, the Jaccard index is strongly affected by dataset size and should be used with caution. The Forbes coefficient (fold change) and tetrachoric correlation are less affected by dataset size, but one should be aware of increased variance for small datasets.

**Availability:** All results on simulated and real data can be inspected and reproduced at https://hyperbrowser.uio.no/sim-measure

## 1 Introduction

A reference genome provides a unified coordinate system to locate where genomic features occur. Genomic tracks [11] indicate the base pair positions on a reference genome where a specific genomic feature is observed. Several large consortia [2, 4, 12] have contributed to the public domain with reference datasets of such genomic features such as cell-specific chromatin states and protein binding regions. Such genomic tracks are often compared against reference collections to identify proteins, chromatin marks, or cell types of interest. These comparisons rely on relevant metrics of similarity (co-occurrence), which should reflect the degree of biological association.

Many of the widely used binary similarity metrics, such as the Jaccard index and the Forbes coefficient (also known as fold change), were initially developed in the context of species presence/absence problems within ecology [5, 9]. Within ecology, there has been a long debate on the properties of different similarity metrics. In the genomics field, however, these metrics have been applied without a critical discussion of their suitedness and typically without providing any reasoning for why a particular metric was chosen. For instance, more than 300 papers on *bioRxiv* between 2010 and 2018, within a limited selection of genomics literature, explicitly mention the Jaccard metric, while the argumentation for such a choice is usually not provided. The Forbes coefficient is also commonly used, but the extent of its usage is harder to quantify. It represents a ratio of observed versus expected co-occurrence and is thus often simply termed *enrichment* or *fold change*. Existing tools for genomic track analysis typically provide only a single metric for the degree of co-occurrence between tracks, without any discussion of the properties and suitedness of this chosen metric. As an example, the widely used BEDTools [15] originally provided only raw overlap counts. Later, the Jaccard index was also added, but without any user guidance on its suitability [14].

In this report, we provide a critical evaluation of the properties of three similarity metrics in the context of genomics: the Jaccard index, the Forbes coefficient, and the tetrachoric correlation. Specifically, we focus on how these metrics may be affected by experimental variability (number of genome-wide annotations per genomic feature) or technical variability (computational assessment of similarity). We introduce two alternative modelling assumptions for how an underlying biological feature may give rise to experimental datasets of varying size (number of base pairs covered by a track). According to each of these assumptions, we carefully assess the robustness of these three commonly used similarity metrics. By connecting the choice of metric to underlying assumptions, we enable the genomics community to make reasoned choices of metrics in particular settings. Furthermore, we perform a large-scale and systematic study on how size variation of experimental datasets affect rankings of co-occurrence according to each of the three similarity metrics.

## 2 Methods

In the present study, we refer to *binary vectors* or *tracks* as sets of occurrences anchored to specific coordinates in a reference genome. Therefore, throughout the study, we interchangeably use the terms similarity and co-occurrence when referring to relations between genomic tracks. A reference track can be defined as a binary indicator *R*, where *r*_*s*_ = 1 if position *s* is covered by an occurrence for *s* ∈ 1, …, *N*, with *N* being the length of a reference genome. We refer to the coverage of a track (sum across the vector) as the *size* of the track. A query track *Q* is a separate track against which reference tracks are analysed.

We introduce two principled models for how an underlying genomic feature may give rise to experimental datasets of varying size. The first model assumes that an experimental dataset of binary events is the imperfect observation of an underlying binary biological process. The second model assumes that an experimental dataset of binary events corresponds to a particular thresholding of an underlying continuous reality.

### 2.1 Model assumption 1: Binary underlying reality

Depending on the context, the underlying reality of a biological process can be considered to be binary (see Supplementary Material for a formal definition). For instance, this could correspond to a transcription factor (TF) either binding to the DNA or not. Under this assumption, a track *R* represents an imperfect observation of the underlying binary reality *Y*. The track *R* could denote the locations of called peaks of a ChIP-seq dataset, where some peaks reflect true binding events (true positives, tp), some peaks do not reflect any true binding event (false positives, fp) and some true binding events remain undetected (false negatives, fn) [17]. A particularly interesting case is when the rate of false positives in *R* is invariant to the dataset size. In the context of a ChIP-seq dataset, this occurs if the false discovery rate has been properly controlled when calling peaks, i.e. when a given proportion of occurrences in *R* reflects true binding sites (the underlying reality *Y*), regardless of the dataset size.

### 2.2 Model assumption 2: Continuous underlying reality

Alternatively, an observed binary dataset can represent the discretisation of an underlying continuous reality. This corresponds to assuming that a TF has a varying propensity to bind and that a dataset represents the set of genomic locations where the binding propensity is above a given threshold.

When assuming a continuous underlying process, the similarity between the underlying continuous features are estimated based on the observed binary occurrences. A particular case is when two observed binary tracks *Q* and *R* represent thresholded instantiations of one underlying bi-normal process [8]. Under this assumption, the correlation of this bi-normal process is a natural similarity metric, and the goal is to estimate the similarity between the latent continuous values based on the observed binary values.

subsectionSimilarity metrics Currently, several metrics have been used to quantify co-occurrence of (similarity between) genomic tracks. As mentioned, two commonly used metrics are the fold enrichment (Forbes coefficient) [5] and the ratio of intersection to union (Jaccard index) [9]. Using set notation, the Forbes coefficient is defined as 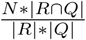, where the notation |*T|* indicates the number of elements of a set *T*. The Jaccard index is defined as 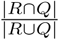. Using the notation in Table 1, the two indexes can be calculated as 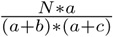 and 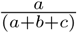 respectively.

**Table 1:** Contingency table

| | $Q = 1$ | $Q = 0$ |
| --- | --- | --- |
| $R = 1$ | a | b |
| $R = 0$ | c | d |

Given our consideration of an underlying continuous reality, a third metric of interest is the tetrachoric correlation, which is a classic measure of correlation [1]. To our knowledge, it has not previously been used to assess similarity (co-occurrence) in a genomic context. The tetrachoric correlation assumes that the two tracks *Q* and *R* are generated by thresholding two underlying continuous processes that are bi-normally distributed, and is defined as the correlation (*ρ*) between these two underlying processes. The thresholds and the correlation *ρ* can be estimated from *Q* and *R* using maximum likelihood techniques [8]. In our study, we have used the *polycor* R package [6, 16] to estimate *ρ*.

### 2.3 Tools

All plots and the information necessary for their reproduction can be found at https://hyperbrowser.uio.no/sim-measure. Plots generated by GHB webtools can be reproduced using the redo-functionality provided by the underlying Galaxy system. Plots generated in R are accompanied by their respective R code and the data files used to generate them. Plots generated by the UCSC Genome Browser are accompanied by URLs and form inputs required to generate similar plots in the current version of UCSC.

## 3 Results

Here, we explore the implications of the two principled models (binary and continuous underlying reality) on three different similarity metrics (Jaccard index, Forbes coefficient, and tetrachoric correlation) using analytical derivations, simulations, and real data. We show that the dataset size may strongly affect the expected similarity value of a particular metric, as well as the uncertainty connected to this value. Further, we investigate the robustness of these metrics by quantifying TF co-occurrence based on a large collection of ChIP-seq peak datasets of variable size [3]. We first derive the theoretical properties of the metrics in each scenario (see Supplementary Material) and present experimental results on simulated and real data.

For the tetrachoric correlation, similarity values are based on methods using maximum likelihood estimation, therefore deriving analytical properties is not feasible.

### 3.1 Theoretical properties of the metrics

#### Model assumption 1: Binary underlying reality

Assuming a binary model, we can show that if *R* is an imperfect observation of *Y* with an expected false discovery rate (FDR) with respect to *Y* of *q*, i.e.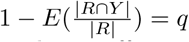, then the expected Forbes coefficient of *R* and *Q* is a function of the Forbes coefficient of *Y* and *Q*, given by:

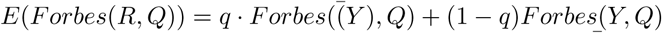 where *Forbes*(*Y, Q*) = *Forbes*(*Y* = 1 and *Q* = 1), while 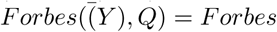 and *Q* = 1). This allows for the Forbes similarity of two tracks *R*^1^ and *R*^2^ to be comparable with respect to *Q*, as long as the FDR of *Y* ^1^ and *Y* ^2^ (*q*) is the same. If the FDR is zero (*q* = 0), *E*(*Forbes*(*R, Q*)) reduces to *Forbes*(*Y, Q*), meaning that the Forbes coefficient of *R* and *Q* provides an unbiased estimate of the Forbes coefficient of *Y* and *Q*. We show this does not hold for the Jaccard index (see derivation given in Supplementary Material), since even for the case of no false positive *E*(*FDR*) = 0, the expected Jaccard index of *R* and *Q* is dependent on the size of *R* (given the subsetting rate *k*):

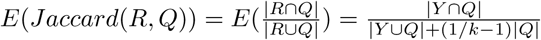

#### Model assumption 2: Continuous underlying reality

When binary vectors *R* are observed, *R* can represent a binary discretisation of an underlying continuous reality *Y* *. Correspondingly, a query track *Q* represents the discretisation of a continuous reality *X*\*. Assuming that such a continuous underlying process exists, one is interested in the similarity between the latent continuous variables *Y* * and *X*\*, given the observed binary vectors *R* and *Q*, as well as the latent discretisation thresholds *t* and *t*\*. Under these conditions, the Forbes coefficient would give a systematic relative bias for small versus large observed tracks, since its values are based on the conditional probabilities *P* (*Q|R*) (details provided in Supplementary Material). An explicit procedure of randomly subsetting occurrences (in order to scale a set of tracks *R* to the same size) would give rise to the same systematic bias. The reason is that the observed occurrences (*R*) of large tracks represent a range of continuous values (*Y* *) that starts at lower values than for smaller tracks (since track size variation is assumed to be due to different thresholds). Thus, the subset of occurrences from a large track *R* represents (on average) lower continuous values of the underlying track (*Y* *). The correlation between *R* and the query is therefore associated with lower continuous values of *X*\* and thus with lower frequency of occurrences at these locations in *Q*. Thus, neither the use of a metric such as Forbes or the use of explicit subsetting procedures are recommended in contexts where the observations represent a thresholded version of a continuous underlying reality. As an example, TF binding regions are typically determined by thresholding a signal that is based on the depth of mapped reads (referred to as ChIP-seq peak calling). If this read depth-based signal correlates with an underlying propensity to bind, then the variation in the dataset size will have the above characteristics.

Under such assumptions, the similarity between the tracks can alternatively be evaluated by the tetrachoric correlation, which assumes that the two tracks *Q* and *R* are generated by thresholding two bi-normally distributed underlying (continuous) processes. The tetrachoric correlation is defined as the correlation (*ρ*) between these two underlying processes (details in Supplementary Material).

### 3.2 Experimental results

To further delineate the behaviour of the three similarity metrics under controlled conditions, we performed a simulation study. Separately for Model assumptions 1 and 2, we generated datasets of varying sizes based on a fixed underlying degree of similarity according to a given model.

#### Simulated data under Model assumption 1

Figure 1 shows how the three considered metrics behave when the datasets are simulated according to Model assumption 1. Datasets of varying sizes, but fixed (expected) false discovery rate of 5%, were generated from the fixed base reference track *Y*, representing the usually unknown underlying track, while a fixed query track *Q* was used. Consistent with the analytical derivations, the Forbes coefficient seems to be less dependent on track size, while the Jaccard index is highly influenced by track size. The Jaccard index drops dramatically when the size of the reference track decreases.The size effect dominates the underlying (biological) signal when large size differences are present. The tetrachoric correlation is also affected by track size, but less than the Jaccard index.

**Figure 1:**
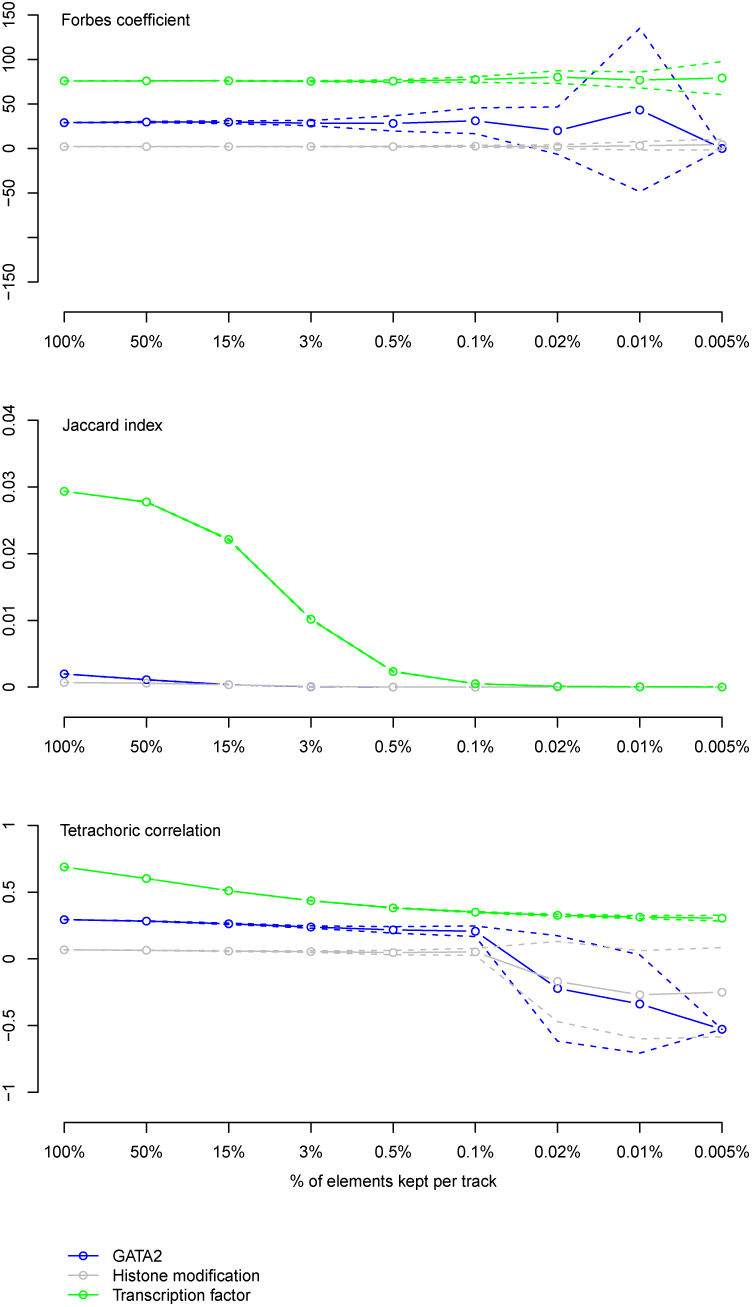
The behaviour of the three considered similarity metrics on simulated datasets with size varying according to Model assumption 1. Top panel: Simulation study showing the mean Forbes coefficient between a fixed query track *Q* and reference tracks *R*s corresponding to varying sizes of a fixed base reference track (*Y*). We set up three simulations based on a GATA1 ChIP-seq track as query track and respectively a GATA2 (blue), a transcription factor (green), and an histone modification (grey) ChIP-seq track as a base reference (details in Methods). The x-axis shows the proportion of occurrences in *R* compared to the reference track *Y*, while the y-axis shows the resulting mean Forbes coefficient against the query track (continuous line) and one standard deviation from the mean Forbes coefficient (dashed lines). The middle and bottom panels show the same for the Jaccard index and the tetrachoric correlation.

As expected, the estimates have high uncertainty for small dataset sizes for the three metrics (Figure 1). The expected Forbes coefficient is constant across dataset sizes and the estimated similarity may take on extreme values (either low or high) for small datasets. A similar tendency is observed for the tetrachoric correlation. Since the Jaccard index is systematically low for small datasets, the effect is less pronounced.

Additionally, the same properties were found when simulating according to 50% FDR (Supplementary Figure S1).

#### Simulated data under Model assumption 2

Figure 2 shows how the three considered metrics behave when datasets are simulated according to Model assumption 2. Datasets of varying sizes were generated by applying different thresholds on a fixed continuous base reference track (*Y*), while a fixed threshold was applied to the underlying continuous underlying reality *X* resulting in the query track *Q*. The tetrachoric correlation (bottom panel) provides a stable and unbiased estimate of the underlying correlation even when applying stringent thresholds (tracks with few occurrences). In contrast, the Forbes coefficient shows a systematic increase with stringent thresholds. The Jaccard index shows a very strong systematic pattern. Interestingly, the threshold level that gives rise to the peak value of the Jaccard index is dependent on the threshold used to construct the fixed query track (details in Supplementary Material).

**Figure 2:**
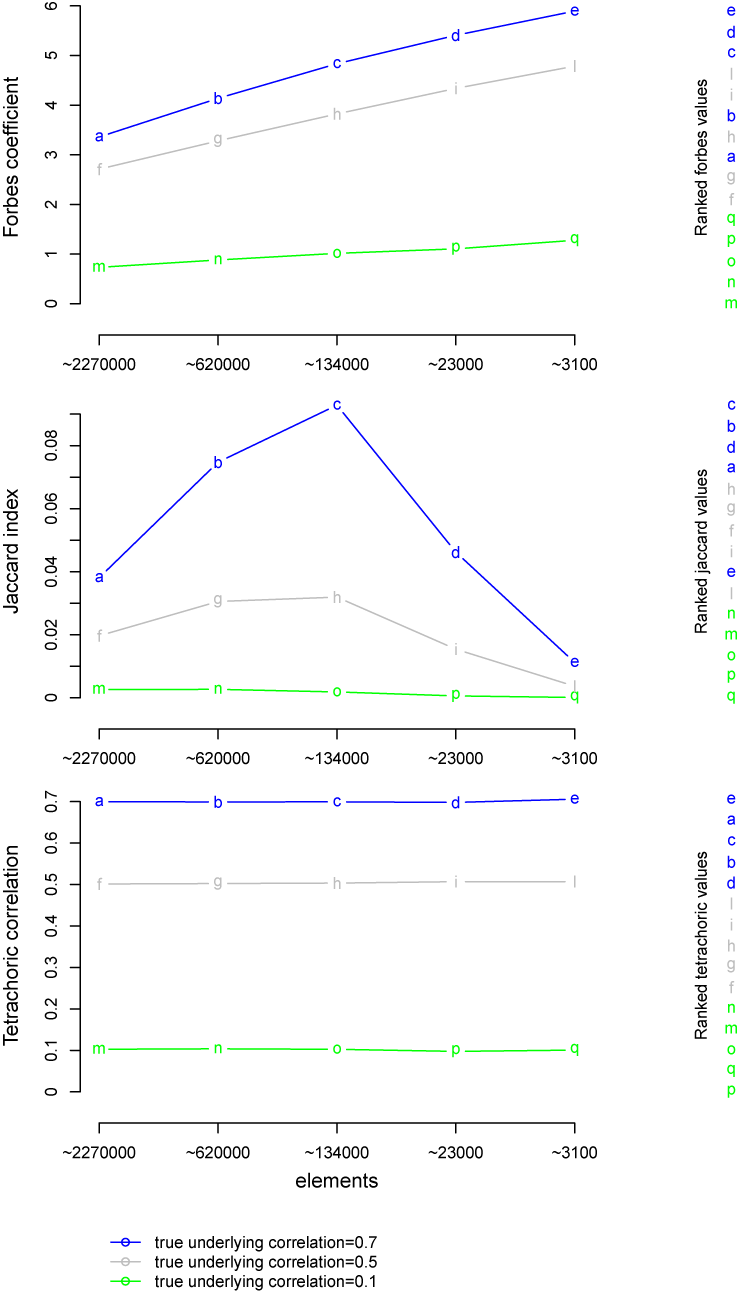
The behaviour of the three considered similarity metrics on simulated datasets with size varying according to Model assumption 2. Top panel: Simulation study showing the Forbes coefficient between a fixed binary query track *Q* and binary reference tracks *R*s corresponding to different thresholding of a fixed, continuous reference track (*Y*). We simulated under three different levels of covariance for the underlying bi-normal process giving rise to *Q* and *Y*, with covariance set to 0.1 (green), 0.5 (grey) and 0.7 (blue). The x-axis shows the number of occurrences in the resulting *R*s after applying different threshold levels for the discretisation of *Y*. The y-axis shows the Forbes coefficient on the discretised *R*s against the fixed query track *Q*. The middle and bottom panels show the same for the Jaccard index and the tetrachoric correlation.

#### Empirical analysis on a large collection of TF ChIP-seq peak datasets

In the simulation studies, we explored how the three considered metrics behave given a particular assumption about the generation of the data. Although consideration of these alternative assumptions helps in choosing an appropriate metric, the appropriate underlying assumption is usually not obvious in real settings. To investigate the behaviour of the different metrics on real data, we performed a large-scale study on ChIP-seq datasets for different TFs. Specifically, we have assessed how the estimated similarity for all pairwise combinations of TF ChIP-seq experiments is affected by the size of the datasets. Figure 3 shows how each of the three considered similarity metrics varies with respect to the size of the experimental datasets. The Jaccard index is consistently low for small datasets and increases with the size of the dataset. The Forbes coefficient and the tetrachoric correlation appear to be less influenced by the track size. The Forbes coefficient shows a downward trend while the tetrachoric correlation shows an upward trend as the track size increases. This suggests that neither Model 1 nor a bi-normal version of Model 2 perfectly models the size variation of the ChIP-seq datasets. Nevertheless, both model assumptions are close and the reality appears to be somewhere in between the two models. For small tracks, the Forbes coefficient shows a higher variability compared to the Jaccard index and to the tetrachoric correlation. The different behaviours with respect to dataset sizes indicate that the choice of metric has a substantial influence on the results. To directly address this question, we compared the ranking of the 36,046 dataset pairs from 269 datasets by co-occurrence when assessed by either the Jaccard index or the Forbes coefficient. Indeed, these two metrics gave highly different rankings. The Spearman correlation for the ranks was 0.54, and 21 of the top 100 dataset pairs for one metric were among the top 100 for the other metric (see Figure 4 for a scatter plot of ranks according to Forbes and Jaccard). Additional figures on the agreement between Jaccard and the tetrachoric correlation, and Forbes and the tetrachoric correlation, are provided as in supplementary Figures S3-S4).

**Figure 3:**
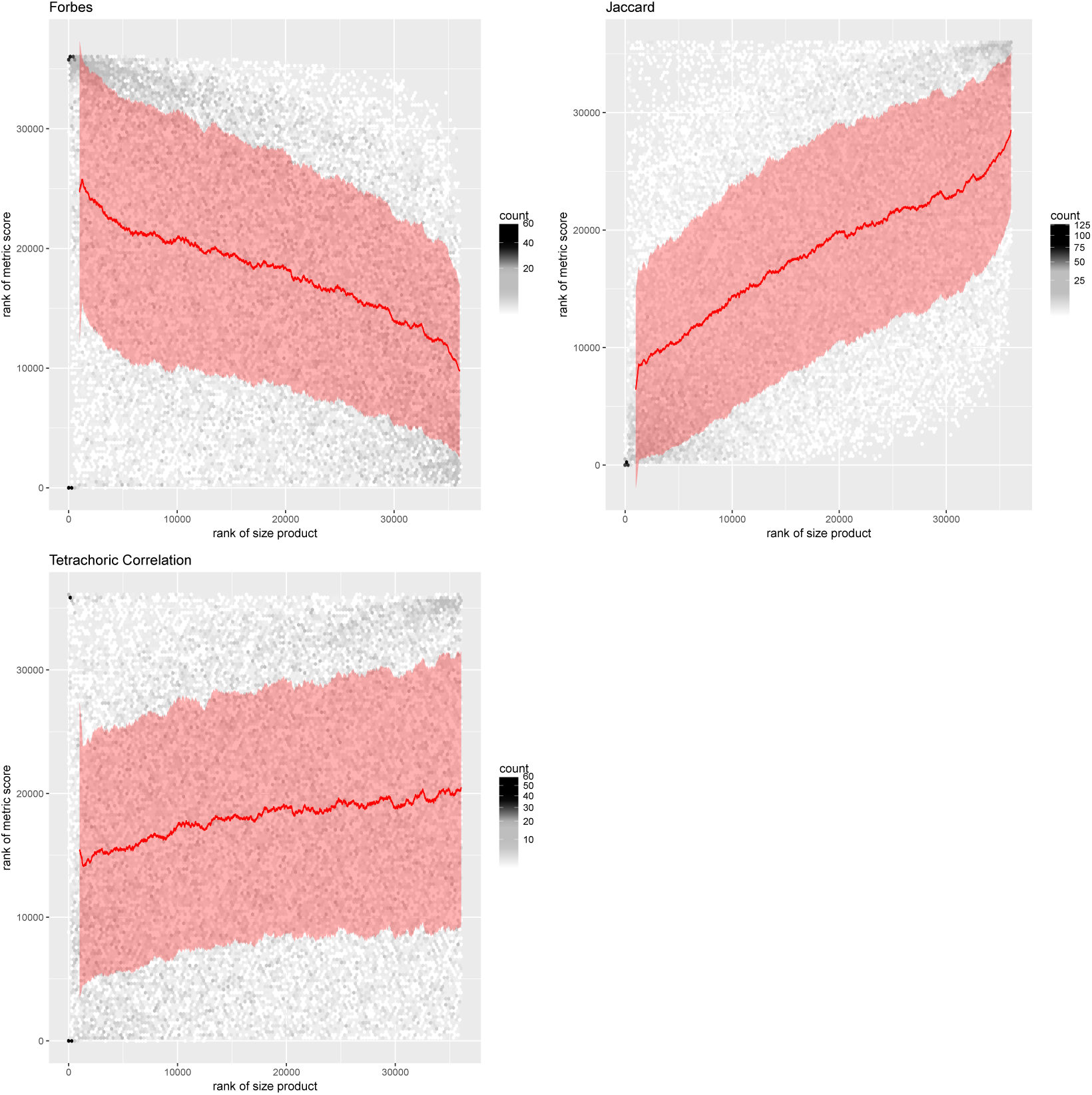
Large-scale empirical analysis of TF co-occurrence assessed by the three considered metrics on ChIP-seq datasets of varying size. Top left panel: Empirical analysis of the Forbes coefficient between 36,046 pairs of experimental (ChIP-seq) datasets for 269 tracks representing binding occurrences for 15 different TFs. The y-axis shows the rank of Forbes coefficient, while the x-axis gives the rank of the product of track sizes for the two experimental datasets compared. The red line and areas show rolling averages +/-rolling standard deviations with a window size of 1000. The top right and bottom panels show the same for the Jaccard index and the tetrachoric correlation, respectively.

**Figure 4:**
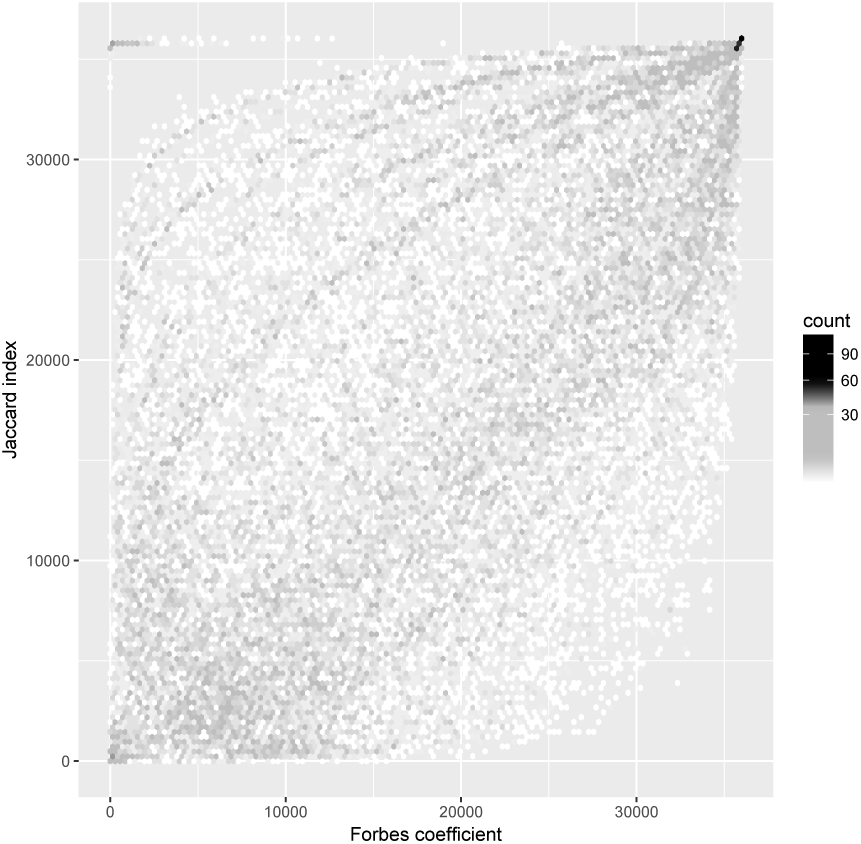
Scatter plot of rankings of large-scale empirical analysis of TF co-occurrence assessed by the Forbes coefficient and Jaccard index on ChIP-seq datasets. The y-axis shows the rank according to the Jaccard index, while the x-axis gives the rank (where a rank equal to one is the least similar) according to the Forbes coefficient for 36,046 pairs of experimental (ChIP-seq) datasets for 15 different TFs.

To further illustrate how Jaccard and Forbes can result in a different conclusion being made when applied on real data, we re-ran a previously published experiment on the co-occurrence of Crohn’s disease-associated variants (GWAS SNPs) and DNase hypersensitive sites (DHSs) [7]. In the original study by Maurano et al., Forbes was used to measure the co-occurrence of the SNPs with DHSs from different cell types, with the conclusion being that in the case of Crohn’s disease, T cell subtypes (therefore, immune cells) show the most-significant GWAS variants in their DHSs (Fig 5, top panel). When we apply Jaccard on the same dataset, we find the complete opposite trend with similarity increasing with DHSs track size (Fig 5, bottom panel).

**Figure 5:**
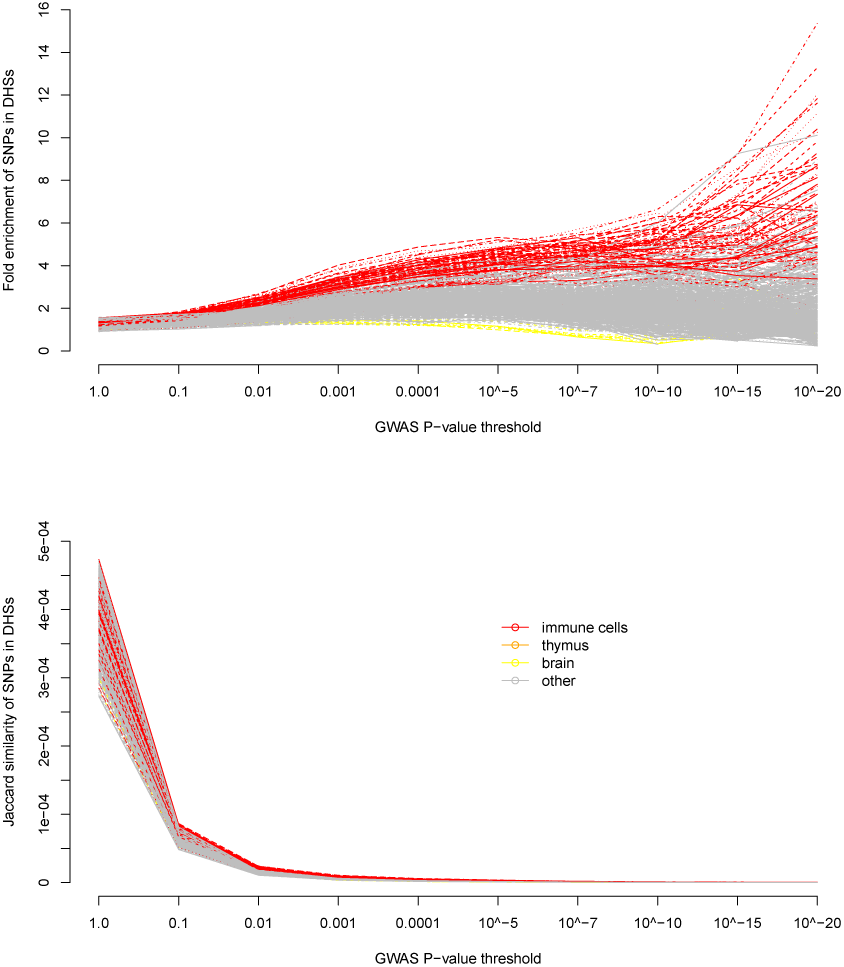
Co-occurrence of Crohn’s disease-associated variants (GWAS SNPs) and DNase hypersensitive sites (DHSs). The top panel, shows the co-occurrence of the SNPs with DHS from different cell types by Forbes (fold enrichment), while the bottom panel shows the same analyses by Jaccard.

## 4 Discussion and conclusion

Here, we have performed an in depth assessment of a central, but often overlooked component of genomic co-occurrence analyses: the similarity metric used to evaluate and compare the degree to which different genomic features co-occur along the genome. Since the number of occurrences may vary widely between experimental datasets, the raw base pair overlap between genomic tracks must be normalised by means of a similarity metric. Two commonly used metrics are the Jaccard index and the Forbes coefficient. Through modelling considerations, we propose that the tetrachoric correlation could also be employed to assess genomic co-occurrence.

We here show that the often overlooked choice of co-occurrence metric may lead to highly disparate rankings of similarity. To better understand what distinguishes the different metrics, we explore their behaviour with respect to datasets size variation according to models assuming either a binary or continuous underlying reality. We show through analytical derivations and simulation that the Forbes coefficient is preferred if the underlying biological feature is assumed to be binary. If an experimental dataset is rather assumed to represent thresholded events from an underlying continuous reality, the tetrachoric correlation may be preferred. A large-scale empirical analysis of TF co-occurrence indicates that the tetrachoric correlation is the least affected by size variation of experimental datasets, while the size variation is highly prone to dominate biological signal if the Jaccard index is used. Nonetheless, one should beware that estimates based on the Forbes coefficient or the tetrachoric correlation have high uncertainty for small tracks. For the tetrachoric correlation, this effect is particularly strong for tracks with mean correlation values close to zero.

While simulated datasets provide full control of the underlying truth, the resulting biases are highly dependent on the models assumed for the simulation. In contrast, the large-scale empirical study shows how the metrics behave under real data generation processes of the considered genomic features (transcription factor binding to DNA). However, for real biological features we only have access to the imperfect experimental observations and have no way of isolating size-associated biases from the true underlying similarity. The strong correlation between the dataset size and the Jaccard index value observed in the TF study could thus, in principle, arise because large datasets mostly represented TFs that were truly more similar (in terms of their true binding locations). Although unlikely, we addressed this possibility in Supplementary Figure S2 by normalising the similarity value for a given pair of experimental datasets against the average similarity for all pairs of experimental datasets for the same pair of TFs. This would thus cancel out any effect due to larger datasets representing more similar TFs. The results after doing this correction were very well in line with those observed in the main figures (for all three metrics). An even subtler point is that large experimental datasets could represent contexts (cell type and cell condition) for which the TFs were systematically behaving more similar to each other, so that even correcting for the pair of TFs would not cancel out variation in true similarity. Since such context would mainly be unspecified, it is not possible to control for. Nonetheless, there is no a priori reason to believe that there would be such systematic associations, and we consider it very unlikely that it could have a strong influence on the results. We also note that the bias we see for the Jaccard index in the empirical study is in line with what we see in the controlled simulations. Additionally, when applying the Forbes coefficient and the Jaccard index to measure the similarity between Crohn’s disease GWAS SNPs and DHSs of specific cell types, as published by [7], we obtain very different results depending on which similarity metric is being used. When assessing colocalization by the Forbes coefficient (as in Maurano et al.), DHS enrichment is increasing with higher SNP significance in the presumably relevant immune cell types (Fig. 5 top panel). In contrast, a corresponding assessment based on the Jaccard index (Fig. 5 bottom panel) shows a strongly decreasing value of colocalization with higher SNP significance. This non intuitive relation arises as an artifact due to the lower track size when focusing only on highly significant SNPs.

Our analysis has specifically focused on how size variation of datasets affects similarity metrics. While there are differences between metrics in terms of the range of values they can take and their ease of interpretation, we see the behaviour with respect to dataset size variation as their defining property. Indeed, in terms of ranking the strength of relations, neither the range nor the interpretation of a metric matters; its sole role is to define an ordering of the discrete points in a three-dimensional space, where the size of the two datasets and their intersection make up the three dimensions. Also when a metric is used as test statistic for Monte Carlo-based statistical testing of genomic colocalization, it is only the resulting ordering that will affect significance [10].

While we have here focused on the setting of genomic co-occurrence, we believe our approach to be equally valuable also for other fields in which similarity metrics are used. This includes questions related to species presence in ecology, which is the field in which several of the considered metrics were originally proposed. The approach could also be useful for other questions in genomics where similarity assessment is central, such as gene set enrichment analysis (GSEA) or various applications of clustering to gene expression or genomic location values. By considering the proposed model assumptions in light of a particular analytical setting, a reasoned choice of a metric can be made. As in our analysis, one should not expect a clear-cut answer, but rather a nuanced conclusion of some metrics appearing more suitable than others.

In conclusion, we find that the choice of similarity metric is an important and often overlooked aspect when analysing genomic co-occurrence. Interestingly, although normalising for size variation is the primary purpose of a similarity metric, we find that the Jaccard index is strongly affected by dataset size, both according to underlying models of size variation and in an empirical investigation. The Forbes coefficient and the tetrachoric correlation show less systematic association with dataset size, and can each be connected to a reasonable underlying model of size variation. Both of these metrics may work satisfactorily in a given context, though one should be aware that estimation uncertainty may lead to extreme values for small datasets. Despite its limited current usage in the field, the tetrachoric correlation indeed shows the most desirable behaviour in our analyses. Ideally, we recommend to explicitly choose between similarity metrics based on a consideration of which of the two underlying models of size variation appear most reasonable for a given biological context. For tools that currently rely on the Jaccard index to quantify genomic co-occurrence, we recommend to include an option of using alternative metrics like the Forbes coefficient and the tetrachoric correlation.

## Acknowledgements

We would like to thank Sveinung Gundersen for his invaluable help with creating the Galaxy page and instance for this specific project.

