## Supplementary material for "Beware the Jaccard: the choice of metric is important and non-trivial in genomic colocalisation analysis"

### 1 Supplementary Methods

#### 1.1 Supplementary on Model 1: Binary underlying model

When the underlying reality of the biological process of interest is discrete (binary vector), we assume that the feature of interest is binary in its nature - i.e. formed by events either occurring or not occurring. In this situation we denote by $R$ an imperfect observation of the underlying binary reality $Y$. If we, for terminology purposes, consider $R$ as a prediction of true binary classes in $Y$, we can define true/false positives/negatives at each location, and derive corresponding values specificity (1-$FPR$) and sensitivity (1-$FNR$) for the false positive and the false negative rates, respectively. A case of particular interest is when the occurrences (ones) of the observed vectors $R$ are assumed to be subsets of the occurrences of the underlying vectors $Y$. This corresponds to a situation where the observation may miss underlying occurrences ($FNR > 0$), but that spurious occurrences are constant ($FPR = c$). A similarity measure on the pairwise similarity between a query track $Q$ and a reference track $R$ should be an unbiased estimate of the same measure applied on $Q$ and $Y$. If not, arbitrary variation in $FNR$ (e.g. due to variability in experimental setup) could strongly influence the estimated similarity.

#### 1.2 Supplementary on Model 2: Continuous underlying model

In certain settings where binary vectors $R$ are observed, it may be natural to assume these to represent binary discretizations of an underlying continuous reality of interest ($Y$). As we are only observing binary values of $R$ resulting from a thresholding of $Y$, we have no information on the underlying distributions for $Y$. As we will show in the following, perhaps contrary to intuition it may still be useful to make assumptions on the underlying distributions of $Y$ and of a threshold $t$ that determines the binary values of $R$ given $Y$. Similarly, we may assume that an observed binary query track $Q$ represents the discretization of an underlying continuous track $X$ (based on a threshold $t^*$). One possibility is to assume that each pair $Q - R$ represent discretizations (based on a threshold pair $t^* - t$) of an underlying latent bi-normal distribution for $X - Y$.

### 2 Supplementary Results

#### 2.1 Theoretical properties of each metric

*Co-occurrence between tracks and conditional probability*

Given a track $R$, we define the corresponding size of the track as the total number of base pairs $s$ covered by a segment for $s \in 1, ..., N$ and $N$ the length of the track. The corresponding coverage probability for a track $R$ is defined as $P(R = 1)$, and can be estimated as $P_R = 1/N * \sum_{s=1}^{N} r_s$. Given a single track $R$ and a query track $Q$, the co-occurrence between the two tracks is the number of base pairs $z$ covered by a segment in both tracks for $z \in 1, ..., N$ which can be represented by a contingency table (Table 1).

Having defined the probability distribution of each track $R$ of the set, we are interested in the co-occurrence probability $P(Q, R) = P(Q = 1, R = 1)$. In case of independence between the query track and the single track, this probability would be equal to the product of the marginal probabilities; $P(Q, R) = P(Q) * P(R)$, while in case of existence of a dependency structure between $Q$ and $R$, the joint probability of

the two tracks is expressed in terms of conditional probabilities
$P(Q,R) = P(Q = 1/R = 1) * P(R = 1) = P(Q/R) * P(R)$. The conditional probability is defined as the probability of an event occurring in one track at a given location given that an event has occurred on the corresponding location in another track, and therefore the probability of $Q = 1$ given that $R = 1$.

*Co-occurrence between tracks using set notation*

We derive approximate expected values for the Jaccard and Forbes metrics for a track $R$ that is a random subsampling of a true $Y$.

$E(\frac{|R \cap Q|}{|R \cup Q|}) \approx \frac{E(|R \cap Q|)}{E(|R \cup Q|)}$

$= \frac{k|Y \cap Q|}{k|Y \cup Q| + (1-k)|Q|}$

$= \frac{|Y \cap Q|}{|Y \cup Q| + (1-k)/k|Q|}$

$= \frac{|Y \cap Q|}{|Y \cup Q| + (1/k-1)|Q|}$

$E(\frac{|R \cap Q|}{|R||Q|}) \approx \frac{E(|R \cap Q|)}{|Q|E(|R|)}$

$\approx \frac{k|Y \cap Q|}{|Q|k|Y|}$

$\approx \frac{|Y \cap Q|}{|Q||Y|}$

This shows that the Forbes metric for $R$ is approximately unbiased for the Forbes metric of $Y$, while the Jaccard metric is not unbiased but is dependent on the subsetting rate $k$.

*Co-occurrence statistic under the binary assumption*

Under the binary assumption of the underlying process, we can count the overlap between the query track ($Q$) and each track $R$, deriving estimates also for the marginal and conditional probabilities.

If a feature vector $R$ represents an occurrence subset of the underlying class variable $Y$, then the conditional probability of $Q = 1$ given that $R = 1$ ($P(Q/R)$) is an unbiased estimate of the conditional probability of $Q = 1$ given that $Y = 1$ ($P(Q/Y)$). This property that the conditional probability is invariant to subsetting can be easily demonstrated.

Table 1: Contingency table

| | $Q = 1$ | $Q = 0$ |
|---|---|---|
| $Y = 1$ | a | b |
| $Y = 0$ | c | d |

Table 2: Contingency table

| | $Q = 1$ | $Q = 0$ |
|---|---|---|
| $R = 1$ | a*k | b*k |
| $R = 0$ | c+(1-k)*a | d+(1-k)*b |

Let us indicate with $k$, $0 < k < 1$ the subsetting rate going from $Y$ to $R$. The subsetting rate is one minus the ratio of ones in $Y$ which becomes zeros in $R$ during the subsampling procedure. The two contingency tables 1-2, show the co-occurrence between the tracks before and after subsampling respectively. By using these two tables, it is easily possible to demonstrate the point being made. The conditional probability of $Q = 1$ given $Y = 1$, is $P(Q/Y) = \frac{P(Q=1,Y=1)}{P(Y=1)} \approx \frac{a}{(a+b)}$, while the conditional probability of $Q = 1$ given $R = 1$, $P(Q/R) = \frac{P(Q=1,R=1)}{P(R=1)} \approx \frac{a*k}{(a*k+b*k)} = \frac{a*k}{(a+b)*k} = \frac{a}{(a+b)}$.

The subsetting assumption, i.e. the assumption that the $FNR$ might be high but that the $FPR$ is constant, may be reasonable in a variety of situations where observations are called with high stringency and limited detection power. In settings where this holds, it is advantageous to use a similarity measure which is invariant to subsetting. Also worth noting is that it appears to be a common practice to explicitly draw a random subset of occurrences down to a shared number of occurrences for each dataset when comparatively analyzing multiple genomic datasets. Such explicit subsetting actually follows from the same assumption that variation in observed track size corresponds to varying $FNR$s with constant $FPR$. Indeed, if using a similarity measure that is unbiased under the subsetting assumption, such explicit subsetting will not affect the expected value and only increase uncertainty of the estimate (and is thus discouraged). If the subsetting assumption does not hold, i.e. if both $FNR$ and $FPR$ are unknown but not constant, then it is not possible to derive a similarity measure for $Q - R$ that is an unbiased estimate of $Q - Y$. However, due to the connection between explicit subsetting and $FNR$, when $FPR$ is not constant, an explicit subsetting procedure suffers from similar limitations as unbiased similarity measures.

Currently, several statistics have been used to quantify overlap between tracks. Two commonly used measures are the fold enrichment (Forbes) and the ratio of intersection to union (Jaccard similarity). The Forbes similarity measure is defined as $\frac{N*|R \cap Q|}{|R|*|Q|}$, where the notation $\mid T \mid$ indicates the number of elements of a set $T$, while the Jaccard similarity measure is defined as $\frac{|R \cap Q|}{|R \cup Q|}$. Using notation in Table 1 , the two indexes can be calculated as $\frac{N*a}{(a+b)*(a+c)}$ and $\frac{a}{(a+b+c)}$ respectively.

The probability notation for the Forbes index can directly be derived from the latter formula by multiplying and dividing both members by N:
$\frac{N*a}{(a+b)*(a+c)} * \frac{N}{N} = \frac{\frac{a}{N}}{\frac{(a+b)}{N} * \frac{(a+c)}{N}} \approx \frac{P(Q=1,R=1)}{P(Q=1)*P(R=1)}$, and can be defined as the ratio between the co-occurrence probability $P(Q,R) = P(Q = 1, R = 1)$ and the product between the marginal probabilities of the two tracks; $P(Q) = P(Q = 1)$ and $P(R) = P(R = 1)$ respectively. So, if the two tracks were independent the co-occurrence probability would be $P(Q,R) = P(Q = 1, R = 1) = P(Q = 1) * P(R = 1)$ and therefore the Forbes coefficient would be equal to 1. As shown by the formula, the Forbes index in case of track independence does not depend on the values of the marginal probabilities and in this scenario is equal to 1. If the independence between the tracks did not hold, then the joint probability between the two tracks $P(Q,R) = P(Q = 1/R = 1) * P(R = 1)$ and the Forbes coefficient would result in $\frac{P(Q=1/R=1)}{P(Q=1)}$. Given the subsetting assumption discussed above, $P(Q/R)$ is an unbiased estimate of $P(Q/Y)$, and thus the Forbes coefficient provides an unbiased estimate of similarity between $Q$ and $Y$ (underlying class variable) by looking at similarity between $Q$ and $R$ (observed vector).

By using the probability notation also for the Jaccard index, $\frac{|R \cap Q|}{|R \cup Q|} = \frac{|R \cap Q|}{|R \cup Q|} * \frac{N}{N} \approx \frac{P(Q=1,R=1)}{P(Q=1 \cup R=1)} = \frac{P(Q=1,R=1)}{P(Q=1)+P(R=1)-P(Q=1,R=1)} = \frac{P(Q=1/R=1)*P(R=1)}{P(Q=1)+P(R=1)-P(Q=1,R=1)}$. In case of independence between the query track $Q$ and the generic track $R$, the conditional probability is equal to the product of the marginal probabilities, and the Jaccard index would result in $\frac{P(Q=1)*P(R=1)}{P(Q=1)+P(R=1)-P(Q=1)*P(R=1)}$. As shown by the formula, even in case of independence between the tracks, the Jaccard index tends to increase when the marginal probabilities increase (the tracks are more expressed) independently by the co-occurrence between them. Moreover, the marginal probability is (obviously) not invariant to occurrence subsetting, which means that $P(R = 1)$ is not an unbiased estimate of $P(Y = 1)$, and therefore the Jaccard index provides biased estimate of similarity between $Q$ and $Y$ by looking at similarity between $Q$ and $R$.

#### 2.2 Co-occurrence statistic under the continuous assumption

Under the assumption of the existence of a continuous underlying process, the question of interest is to determine the similarity between the latent continuous variables $X$ and $Y$ given the observed binary vectors ($Q$) and $R$, and latent discretization thresholds $t^*$ and $t$. Given some specific values of the latent variable $Y$, different values of the latent threshold $t$ leads to a varying number of ones in the binary vector $R$. If one defines $W$ as a vector of internal ranks of the corresponding values of $Y$, variation of threshold $t$ can be considered as variation of how many of the top ranks in $W$ are included as values one in track $R$. One may assume that all variation in number of occurrences in a track $R$ is due to differences in values $t$, so that $Y$ e.g. follow a similarly scaled normal marginal distribution. So, the similarity measure between the observed query track $Q$ and the generic track $R$ should ideally be an unbiased estimate of the true similarity between the underlying processes $X$ and $Y$, and therefore invariant to thresholding. As the underlying processes $X$ and $Y$ are continuous, the similarity could e.g. be defined as the Pearson correlation (which is equal to the covariance divided by the product of the standard deviation of the two variables if $X - Y$ is considered a bi-normal distribution.

Note that given $t > z$ leading to binary discretizations $R, Z$ for the same $Y$, the occurrences in $R$ will be a subset of those in $Z$. However, in contrast to the subsetting discussed in the section on discrete latent tracks, $R$ is here not a random subset of $Z$, but rather represents a systematic selection of the most highly ranked values of the latent distribution $Y$. Given a positive correlation between $X$ and $Y$, the indexes of the most highly ranked values of $Y$ will also (by definition, given positive covariation) tend to contain higher-ranked values in $X$. This again corresponds to indexes more likely to contain ones in $Q$. Thus, a smaller track $R$ is associated with higher values in $Y$ (due to higher threshold $t$), which is a again associated with higher values in $X$ (given positive correlation), which is finally associated with the value of $Q$ at that index to be one. Thus, $P(Q|R)$ is not invariant to the number of occurrences in R. On the contrary, given a fixed underlying positive correlation for $X - Y$ and a track $R^*$ having more occurrences than $R^-$, $P(Q|R^*)$ would be expected to be lower than $P(Q|R^-)$. In other words, given the assumption of thresholded latent bi-normal distribution for $X - Y$, a measure based on $P(Q|R)$ (such as Forbes) would give a systematic relative bias for small versus large observed tracks. Similarly, an explicit procedure of subsetting occurrences in order to scale a generic track $R$ of a set, to the same size would lead to the (randomly) selected occurrences corresponding to indexes of higher latent ranks, and thus to a systematic relative bias for tracks that were originally small versus large. Thus, neither the use of measures such as Forbes or the use of explicit subsetting procedures are advisable in contexts where it is natural to assume that the observations represents a thresholded version of a continuous underlying reality.

Under such assumptions, the similarity between the tracks can instead be evaluated by e.g. the tetrachoric correlation. The tetrachoric correlation assumes that the two tracks $Q$ and $R$ are generated by thresholding two underlying continuous processes bi-normally distributed [**7**], and is defined as the correlation ($\rho$) between the two underlying processes. In general, the thresholds can be different for $Q$ and $R$, and together with the correlation $\rho$, can be estimated using maximum likelihood techniques. In our study, we have used the R-package *polycor* [**6**] to estimate $\rho$.

#### 3 Supplementary Figures

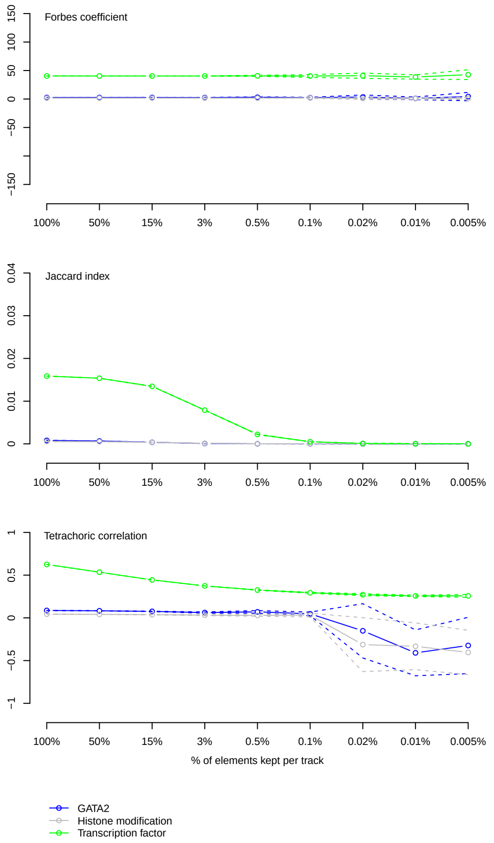


Figure 1: The behaviour of the three considered similarity metrics on simulated datasets with size varying according to Model 1 and stable FDR of 50%. Top panel: Simulation study showing the Forbes coefficient between a fixed query track $Q$ and reference tracks $R$s corresponding to varying different sizes of a fixed base reference track ($Y$). We set up three simulations based on a GATA1 chip-seq track as query track and respectively a GATA2 (blue), a transcription factor (green), and an histone modification (grey) ChIP-seq track as base reference (details in Methods). The x-axis shows the proportion of occurrences in $R$ compared to the reference track $Y$, while the y-axis shows the resulting mean Forbes coefficient against the query track (continuous line) and one standard deviation from the mean Forbes coefficient (dashed lines). The middle and bottom panels show the same for the Jaccard index and the tetrachoric correlation, respectively.

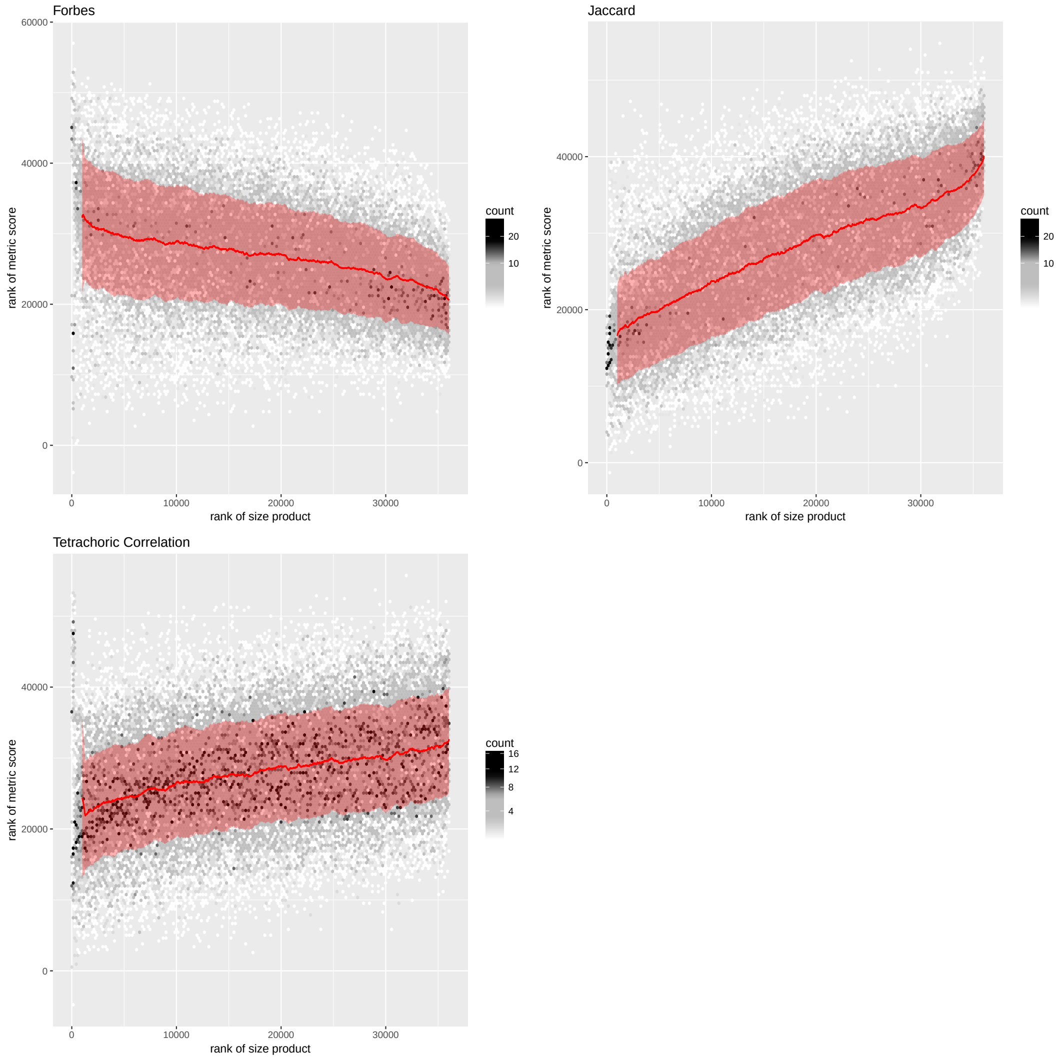


Figure 2: Large-scale empirical analysis of TF co-occurrence assessed by the three considered metrics on ChIP-seq datasets of varying size by subtracting the group effect. Top left panel: Empirical analysis of the Forbes coefficient between 38562 pairs of experimental (ChIP-seq) datasets for 15 different TFs. The y-axis shows the rank of Forbes coefficient after subtracting for group effect, while the x-axis gives the rank of the product of track sizes for the two experimental datasets compared. The red line and areas shows rolling averages +/- rolling standard deviations with a window size of 1000 for the track size rank index. The top right and bottom panels show the same for the Jaccard index and the tetrachoric correlation.

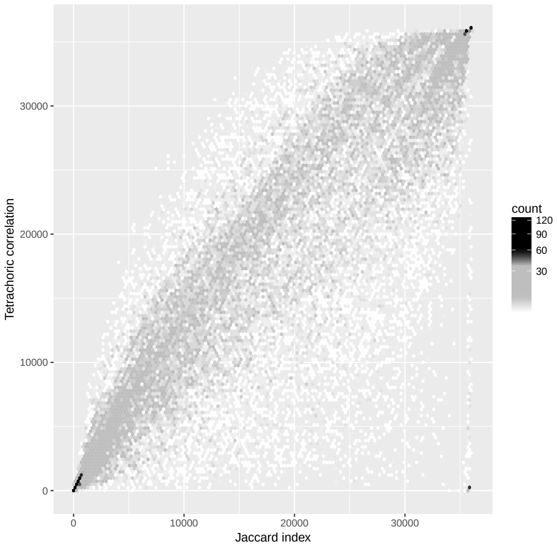


Figure 3: Scatter plot of rankings of large-scale empirical analysis of TF co-occurrence assessed by the Jaccard index and tetrachoric correlation on ChIP-seq datasets. The y-axis shows the rank of tetrachoric correlation, while the x-axis gives the rank (where a rank equal to one is the least similar) of the Jaccard index between 38562 pairs of experimental (ChIP-seq) datasets for 15 different TFs.

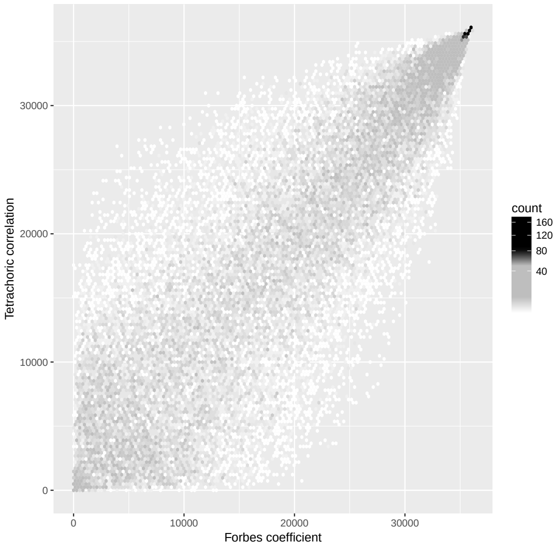


Figure 4: Scatter plot of rankings of large-scale empirical analysis of TF co-occurrence assessed by the Forbes Coefficient and tetrachoric correlation on ChIP-seq datasets. The y-axis shows the rank of Forbes Coefficient, while the x-axis gives the rank (where a rank equal to one is the least similar) of the tetrachoric correlation between 38562 pairs of experimental (ChIP-seq) datasets for 15 different TFs.
